# Hippocampal-cortical encoding activity predicts the precision of episodic memory

**DOI:** 10.1101/2020.11.10.376814

**Authors:** Saana M. Korkki, Franziska R. Richter, Jon S. Simons

## Abstract

Our recollections of past experiences can vary both in the number of specific event details accessible from memory and the precision with which such details are reconstructed. Prior neuroimaging evidence suggests the success and precision of episodic recollection to rely on distinct neural substrates during memory *retrieval*. In contrast, the specific *encoding* mechanisms supporting later memory precision, and whether they differ from those underlying successful memory formation in general, are currently unknown. Here, we combined continuous measures of memory retrieval with model-based analyses of behavioural and neuroimaging data to tease apart the encoding correlates of successful memory formation and mnemonic precision. In the MRI scanner, participants encoded object-scene displays, and later reconstructed features of studied objects using a continuous scale. We observed overlapping encoding activity in inferior prefrontal and posterior perceptual regions to predict both which object features were later remembered versus forgotten, and the precision with which they were reconstructed from memory. In contrast, hippocampal encoding activity significantly predicted the precision, but not overall success, of subsequent memory retrieval. The current results identify a hippocampal-cortical encoding basis for episodic memory precision, and suggest a contribution of shared cortical encoding mechanisms to the formation of both accessible and precise memory representations.

## Introduction

Our memories are not an exact reproduction of the past, but can range from high-fidelity, precise, reconstructions of previous experiences to less accurate, lower-resolution, representations (Brady, Konkle, Gill, Oliva, & Alvarez, 2013; Harlow & Donaldson, 2013). Behavioural evidence suggests such variation in mnemonic precision to be distinguishable from the general success of retrieval (Harlow & Yonelinas, 2016; Richter, Cooper, Bays, & Simons, 2016, but see Schurgin, Wixted, & Brady, 2020), with these two memory components found to be differentially sensitive to various experimental manipulations (e.g., Berens, Richards, & Horner, 2020; Sun et al., 2017; Sutterer & Awh, 2016; Xie & Zhang, 2017), as well as to memory impairments in distinct populations (Cooper et al., 2017; Korkki, Richter, Jeyarathnarajah, & Simons, 2020; Nilakantan, Bridge, VanHaerents, & Voss, 2018). Furthermore, the success and precision of episodic recollection appear to recruit at least partially separable neural mechanisms (Nilakantan, Bridge, Gagnon, VanHaerents, & Voss, 2017; Richter et al., 2016; Ritchey & Cooper, 2020). In particular, during memory *retrieval*, activity in distinct regions of the posterior-medial network has been shown to support each aspect of memory performance (Richter et al., 2016). However, despite increased interest in the neural basis of mnemonic precision (e.g., Cooper & Ritchey, 2019; Montchal et al., 2019; Richter et al., 2016; Stevenson et al., 2018), the associated encoding mechanisms, and whether they differ from those supporting successful encoding in general, remain unresolved.

Successful episodic memory formation is typically associated with activity increases in a network of medial temporal, lateral prefrontal, and posterior perceptual regions (Kim, 2011; Spaniol et al., 2009). The hippocampus receives input from content-specific perceptual regions, and is thought to bind disparate event features into a coherent memory representation (Cooper & Ritchey, 2020; Davachi, 2006; Paller & Wagner, 2002; Ranganath, 2010), and allow for the storage of similar experiences in an orthogonalised, or non-overlapping, manner (Norman & O’Reilly, 2003; O’Reilly & McClelland, 1994). Lateral prefrontal regions, on the other hand, are involved in the strategic and controlled encoding of information into memory via selection, elaboration and integration of goal-relevant information (Blumenfeld & Ranganath, 2007; Simons & Spiers, 2003). The specific neural substrates supporting successful memory formation have been found to exhibit process-specificity, varying for instance according to the depth of stimulus processing engaged in at encoding (Fletcher et al., 2003; Otten et al., 2001), and the type of retrieval process later recruited (Ranganath et al., 2004; Staresina & Davachi, 2006). Moreover, encoding correlates appear sensitive to more subtle differences in the quality of retained representations, including their objective amount of detail (Cooper & Ritchey, 2020; Qin et al., 2011), and subjective strength or vividness (Kensinger, Addis, & Atapattu, 2011; Qin et al., 2011). However, while beginning to elucidate the encoding mechanisms underlying variation in more qualitative aspects of later retrieval, prior studies have focused on the amount details (e.g., number of specific event attributes), or relied on participants’ subjective reports, leaving the specific encoding substrates underpinning more fine-grained variations in objective memory fidelity unclear.

It is possible that, in addition to relying on distinct brain regions during *retrieval* (Richter et al., 2016), the success and precision of episodic recollection may be supported by at least partly separable neural mechanisms during memory *encoding*. For instance, the successful retrieval of information from memory may depend on the strength of an association between a retrieval cue and the target memory, thus drawing in particular on associative encoding processes supported by the hippocampus and the prefrontal cortex (Blumenfeld & Ranganath, 2007; Davachi, 2006). In contrast, the precision with which specific mnemonic features can be reconstructed from memory may closely relate to the fidelity of stimulus encoding in posterior perceptual regions (Emrich et al., 2013), and/or to hippocampal function supporting the formation of distinct and detailed memory traces that can be later reconstructed with high precision (Moscovitch et al., 2016; Yonelinas, 2013). Alternatively, it is possible that, contrary to dissociable neural substrates observed during retrieval (Richter et al., 2016), the successful and precise encoding of information into memory may rely on similar neural mechanisms that perhaps act to increase the strength of the memory more generally, rendering it both accessible and precise at retrieval (Schurgin et al., 2020).

In the current study, we employed continuous measures of memory retrieval and model-based analyses of behavioural and neuroimaging data to elucidate the encoding substrates of mnemonic precision. In the MRI scanner, participants encoded visual stimulus displays depicting an object overlaid on a scene background. The location and colour of the objects were drawn from circular spaces, and at retrieval, participants recreated these attributes of the studied items using a continuous response dial. This approach allowed us to segregate encoding activity supporting later successful memory retrieval from that supporting subsequent mnemonic precision in a manner not afforded by more typical categorical measures of retrieval performance (e.g., old/new, remember/know), thus providing novel insight into the encoding mechanisms supporting the acquisition of high-fidelity episodic memories.

## Methods

### Participants

Twenty-one young adults (18-29 years old) participated in the current experiment. All participants were right-handed, native English-speakers, had normal or corrected-to-normal vision, no colour blindness, and no current or historical diagnosis of any neurological, psychiatric, or developmental disorder, or learning difficulty. Participants indicated no current use of any psychoactive medication, and no medical or other contradictions to MRI scanning. One participant was excluded from all analyses due to excessive movement (> 4mm) in the scanner, leaving 20 participants to contribute to the present analyses (8 male, 12 female; mean age 22.15 years, *SD*: 3.10). The participants were recruited via the University of Cambridge Psychology Department Sona volunteer recruitment system (Sona Systems, Ltd) and community advertisements, and were reimbursed with £30 for their participation. All participants gave written informed consent in a manner approved by the Cambridge Psychology Research Ethics Committee.

### Materials

Stimuli for the memory task comprised 180 images of outdoor scenes and 180 images of distinct everyday objects. The images were obtained from existing stimuli sets (scenes: Richter et al., 2016; objects: Brady et al., 2013) and Google image search. Each object image was randomly paired with a scene image to generate a total of 180 trial-unique study displays (size 750 x 750 pixels). Across the study displays, we varied the appearance of two object features: colour and location. For each display, object colour and location were randomly selected from circular parameter spaces (0-360°) (cf. Cooper et al., 2017; Richter et al., 2016) (see Figure 1). All participants viewed the same study displays in a randomized order.

**Figure 1.**
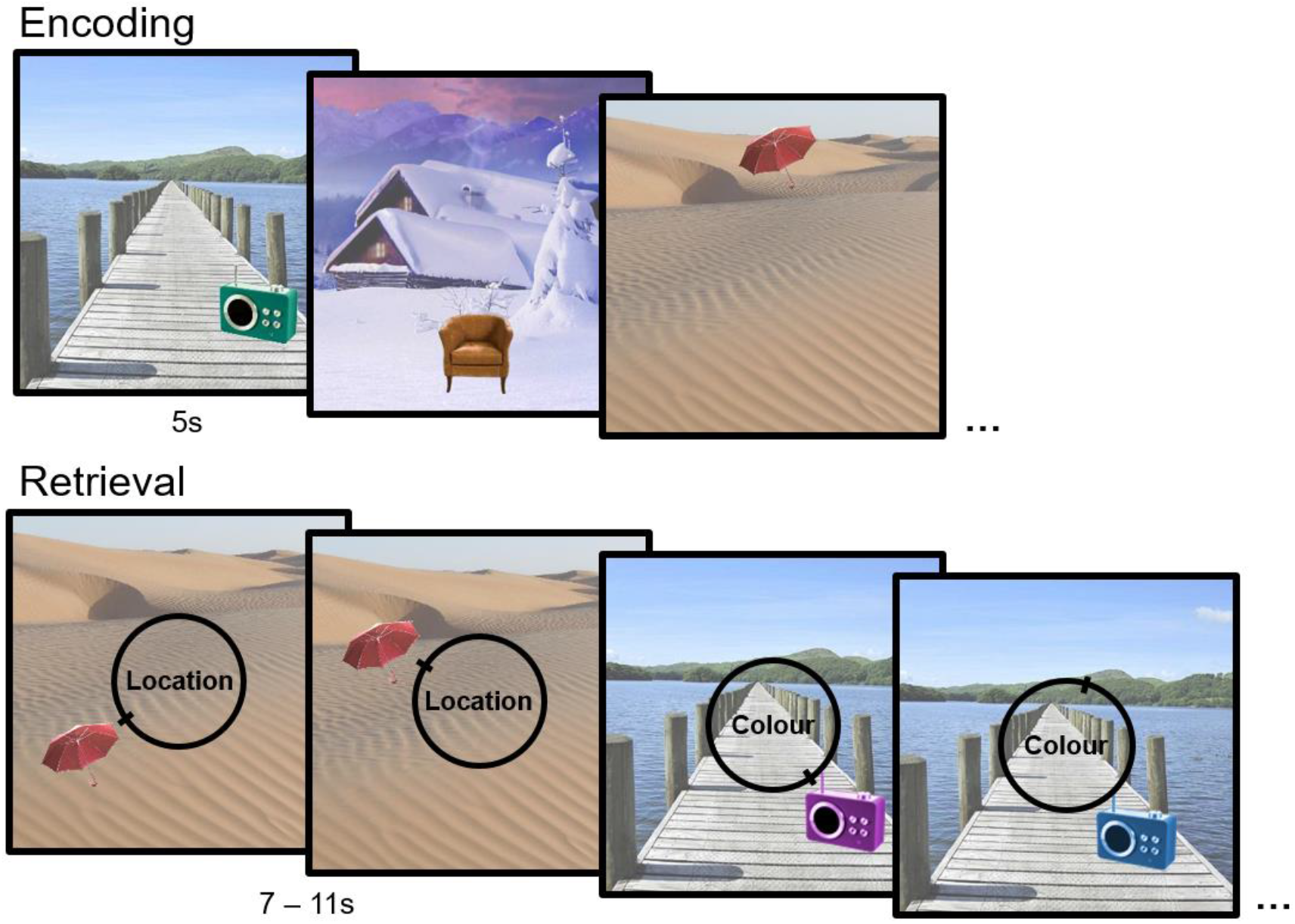
Example study and test trials of the memory task. At study, participants viewed stimuli displays consisting of one object overlaid on a scene background (stimulus duration: 5s). The location and colour of the objects at study were randomly chosen from circular parameter spaces (0-360°). At test, participants recreated *either* the location *or* the colour of each studied object using a 360-degree continuous response dial, allowing for a fine-grained assessment of memory fidelity.

### Design and procedure

Prior to the scan, participants read the instructions and undertook practice trials of the memory task. The task was adapted from Richter et al. (2016) to optimize the investigation of encoding-related effects, and comprised 9 study-test blocks completed over 9 functional runs (one study and one test phase per run). At study, participants sequentially viewed 20 object-background displays (stimulus duration: 5s), and were instructed to try and memorise the appearance of each display, including the location and colour of the object. The study phase was followed by a 10s delay during which a “Get Ready” message was presented on a black screen. Following this delay, participants were asked to reconstruct *either* the location *or* the colour of each object viewed in the preceding study phase (one feature question per each encoding trial, total of 20 retrieval trials per block). At retrieval, the test object reappeared on its associated background with a central cue word “Location” or “Colour” indicating the type of feature tested on that trial. The initial appearance of the tested feature was randomly selected from a circular parameter space (0-360°), while the appearance of the untested feature remained unchanged from study to test. In other words, for location trials, the test object reappeared in its original colour, but in a randomly selected location, whereas for colour trials the test object reappeared in its original location but in a randomly chosen colour. Participants were asked to recreate the object’s original features as accurately as they could by moving a slider around a 360-degree response dial using their middle and index finger on a button box, and were able to confirm their answer by pressing a third key with their thumb. The retrieval phase was self-paced with the constraint of a minimum trial length of 7s and a maximum response time of 11s. Participants on average produced response times that were well under this limit (*M:* 5.64s, *SD:* 0.68s), and the percentage of trials where response selection was not confirmed in time was very low (*M*: 1.36%, *SD*: 1.77%). Note that if a participant failed to confirm their answer within 11s, their last position on the response wheel was recorded as their answer for that trial.

Participants completed 90 location and 90 colour trials in total (10 trials of each type per task block). The type of feature tested for each object was randomised across displays, but constant across participants. To ensure that memory was tested for feature values spanning the entire circular space, this randomisation was conducted with a constraint of roughly equal number (i.e., 20-25) of target feature values sampled from each quadrant around the circular space for both the location and colour condition. The order of study and test displays was randomised across participants with the constraint of no more than four encoding or retrieval trials in a row for which the same type of feature was tested. Study and test trials were separated by a fixation cross with jittered duration between 0.4s and 2.4s (mean ISI duration: 1s) following an approximate Poisson distribution. After the first five of the nine task blocks, participants were given a 10-minute break from the memory task in the scanner, during which a diffusion-weighted structural scan was acquired (analysis of diffusion-weighted data not reported here).

### Behavioural analysis

For each trial, we calculated participants’ retrieval error as the angular difference between their response value and the target feature value (0 ± 180°). To distinguish the likelihood of successful memory retrieval from the precision of the retrieved information, we fitted a two-component mixture model (Bays et al., 2009; Zhang & Luck, 2008) to each participant’s retrieval error data using maximum likelihood estimation (code available at: https://www.paulbays.com/code/JV10/index.php). This mixture model has previously been shown to characterize long-term memory performance in similar tasks (e.g., Brady et al., 2013; Korkki et al., 2020; Richter et al., 2016), and has been employed to gain insights about the neural basis of the precision of episodic recollection (Cooper & Ritchey, 2019; Richter et al., 2016; Stevenson et al., 2018). The model assumes that two distinct sources of error contribute to participants’ retrieval performance across trials: variability, that is, noise, in successful retrieval of target features, and the presence of random guess responses where memory retrieval has failed to bring any diagnostic information about the target to mind. These two sources of error are modelled by a von Mises distribution (circular equivalent of a Gaussian distribution) centred at a mean error of zero degrees from the target value, with a concentration, *K*, and a circular uniform distribution with a probability, *pU*, respectively. Precision of memory retrieval can be estimated as the concentration parameter (*K*, higher values reflect higher precision) of the target von Mises distribution, and the likelihood of successful memory retrieval (*pT*) as the probability of responses stemming from the target von Mises distribution (*pT* = 1 – *pU*). Consistent with prior studies (Korkki et al., 2020; Richter et al., 2016), this two-component model was found to fit the current data better than an alternative one-component model where participants’ responses were assumed to stem from a von Mises distribution around the target feature value only (mean Bayesian Information Criterion (BIC) for the one-component model: 386.10; mean BIC for the two-component model: 317.62).

### MRI acquisition

MRI scanning took place at the University of Cambridge Medical Research Council Cognition and Brain Sciences Unit using a 3T Siemens Tim Trio scanner (Siemens, Germany) with a 32-channel head coil. For each participant, a whole brain structural image was acquired using a T1-weighted 3D magnetization prepared rapid gradient echo (MPRAGE) sequence (repetition time (TR): 2.25s, echo time (TE): 3ms, flip angle = 9°, field of view (FOV): 256 x 256 x 192mm, resolution: 1mm isotropic, GRAPPA acceleration factor 2). Functional data were acquired over 9 runs each comprising one task block (one encoding and one retrieval phase), using a single-shot echoplanar imaging (EPI) sequence (TR: 2s, TE: 30ms, flip angle° = 78, FOV: 192 x 192mm, resolution: 3mm isotropic). Each volume consisted of 32 sequential oblique-axial slices (interslice gap: 0.75mm) acquired parallel to the anterior commissure – posterior commissure transverse plane. Across the participants, the mean number of volumes acquired per functional run was 161.60 (*SD*: 9.63). The scanning protocol further included a diffusion-weighted structural scan that was acquired after the first five functional runs (not analysed here).

### fMRI preprocessing

Data preprocessing and analysis was performed with Statistical Parametric Mapping (SPM) 12 (http://www.fil.ion.ucl.ac.uk/spm/) implemented in MATLAB R2016a. The first five volumes of each functional run were discarded to allow for T1 equilibration. Furthermore, any additional volumes collected after each task block had finished were discarded for each participant so that the last volume of each run corresponded to a time point of ∼2s after the last fixation cross. The functional images were spatially realigned to the mean image to correct for head motion and temporally interpolated to the middle slice to correct for differences in slice acquisition time. The anatomical image was coregistered to the mean EPI image, bias-corrected and segmented into different tissue classes (grey matter, white matter, cerebrospinal fluid). These segmentations were used to create a study-specific structural template image using the DARTEL (Diffeomorphic Anatomical Registration Through Exponentiated Lie Algebra) toolbox (Ashburner, 2007). The functional data was normalized to MNI space using DARTEL and spatially smoothed with an isotropic 8mm full-width at half-maximum (FWHF) Gaussian kernel.

### fMRI analysis

In order to obtain trial-specific estimates of the success and precision of memory retrieval for the fMRI analyses, we fitted the two-component mixture model (von Mises + uniform distribution) to all retrieval errors across all participants (3600 trials in total), and calculated the cut-off point at which the probability of participants’ responses stemming from the target von Mises distribution was less than 5%, beyond which responses likely reflected guessing (cf., Cooper et al., 2017; Richter et al., 2016). This cut-off point was established as +/-51 degrees in the current data, and was then used to classify each encoding trial as successful (absolute subsequent retrieval error ≤ 51°) or unsuccessful (absolute subsequent retrieval error > 51 degrees). Consistent with prior studies (Cooper et al., 2017; Cooper & Ritchey, 2019, 2020; Richter et al., 2016), we used an across-participant model-derived cut-off to ensure that responses of the same error magnitude were consistently classified as successful or unsuccessful across individuals, as well as to avoid any bias in the error cut-offs due to differences in individual model fits. We further note that using feature-specific cut-offs, rather than the threshold estimated across all retrieval trials, did not change the significance of our results. For trials classified as successfully encoded, a trial-specific measure of memory precision was further calculated as 180 – participant’s absolute retrieval error on that trial so that higher values (smaller error) reflected higher precision (range: 129 – 180°) (cf., Cooper et al., 2017; Richter et al., 2016).

For each participant, a first level General Linear Model (GLM) was constructed containing three regressors corresponding to each event of interest (successful location encoding, successful colour encoding, unsuccessful encoding), and a fourth regressor modelling the retrieval trials. For the successful encoding trials, the trial-specific estimates of memory precision were included as parametric modulators comprising two additional regressors in the model. The precision parametric modulators were rescaled to range between 0 and 1 to facilitate the direct comparison of success and precision-related effects, and mean centred for each participant. Neural activity was modelled with a boxcar function convolved with the canonical hemodynamic response function (HRF), with a duration of 5s for the encoding trials and a variable duration (7s-11s) for the retrieval trials, capturing the duration of the study and test displays, respectively. Six participant-specific movement parameters estimated during realignment (3 rigid-body translations, 3 rotations) were further included as covariates in the first level model to capture any residual movement-related artefacts. Due to the small number of guessing trials in each functional run, data from all functional runs were concatenated for each participant, and 9 constant block regressors included as additional covariates. Autocorrelation in the data was estimated with an AR(1) model and a temporal high pass filter with a 1/128 Hz cut-off was used to eliminate low frequency noise. First level subject-specific parameter estimates were submitted to second level random effects analyses.

### Contrasts

The contrasts for the fMRI analyses focused on identifying regions where encoding activity positively predicted the subsequent success and/or precision of episodic memory retrieval (i.e., increases in BOLD signal for successful encoding, or higher memory precision). To examine encoding activity associated with the subsequent success of memory retrieval, we contrasted encoding trials for which memory retrieval subsequently succeeded against trials for which memory retrieval subsequently failed (*subsequent retrieval success effects*). To identify encoding activity predicting the later precision of memory retrieval, positive correlations between BOLD signal and the precision parametric modulator were examined (i.e., linear relationship between BOLD signal and precision parametric modulator; *subsequent precision effects*). We further assessed the overlap between subsequent success and subsequent precision effects using conjunction analyses. Conjunction analyses were conducted testing against the conjunction null hypothesis to ensure that regions identified in this analysis displayed reliable encoding activity associated with each individual contrast, i.e., both subsequent success and subsequent precision of memory retrieval (see Nichols et al., 2005).

Due to a relatively low number of guess trials per feature condition for some individuals, it was not possible to investigate feature-specific subsequent success effects. Furthermore, analysis of feature-specific subsequent precision effects did not yield any significant differences across the ROIs (*p*s > .208), or the whole brain (*p*s > .303). Thus, our analyses focused on examining BOLD activity predicting the subsequent success and precision of memory retrieval across the features conditions, consistent with the approach taken in previous studies employing a similar paradigm (Richter et al., 2016; Cooper et al., 2017).

### Regions of interest

The main analyses focused on a small number of a priori regions of interest (ROIs) implicated by meta-analytic evidence in supporting the successful formation of episodic memories for visual information (Kim, 2011; Spaniol et al., 2009). Specifically, the ROIs included the hippocampus (HC), the inferior frontal gyrus (IFG) and the fusiform gyrus (FFG). Given evidence for greater consistency of subsequent memory effects in the left hemisphere (Spaniol et al., 2009), left-lateralized ROIs were used, each comprising the left anatomical region as defined by the Automated Anatomical Labelling (AAL) atlas. Statistical significance within each anatomical ROI was assessed using small-volume correction with a peak-level familywise error (FWE) corrected (based on random field theory) threshold of *p* < .05, correcting for the number of voxels in each ROI. In addition to the ROI analyses, we sought to identify any additional brain regions displaying a relationship between encoding activity and the subsequent success and/or precision of memory retrieval in exploratory whole brain analyses conducted at a whole brain FWE-corrected threshold of *p* < .05, minimum extent of 5 contiguous voxels.

## Results

### Behavioural results

For each trial, we calculated participants’ retrieval error as the angular difference between their response value and the target feature value (0±180°) (see Figure 2A). Across participants and feature conditions, overall task performance, as measured by the mean absolute retrieval error, was 30.43° (*SD*: 15.04°), with significantly higher mean absolute error in the colour (*M*: 34.48°, *SD*: 15.90°) in comparison to the location condition (*M*: 26.37°, *SD*: 15.80°), *t*(19) = 3.63, *p* = .002, *d* = 0.81. To further decompose the specific sources of error contributing to participants’ overall performance, we fitted the two component mixture model (von Mises + uniform distribution) to each individual participant’s retrieval error data using maximum likelihood estimation (Bays et al., 2009). The mean model-estimated probability of successful memory retrieval, defined as the probability of responses stemming from a von Mises distribution centred at the target feature value (*pT*), was 0.73 (*SD*: 0.18) across participants and feature conditions (see Figure 2B). The mean model-estimated precision of memory retrieval, estimated as the concentration parameter, *K*, of the target von Mises distribution, was 16.79 (*SD*: 7.92) across participants and feature conditions (see Figure 2B) (note that this value of *K* is comparable to an *SD* of approximately 14.20°). Mean memory precision (*K*) was significantly higher in the location (*M*: 34.65, *SD*: 27.24) in comparison to the colour condition (*M*: 10.94, *SD*: 7.15), *t*(19) = 4.04, *p* = 001, *d* = 0.90, whereas mean probability of successful memory retrieval (*pT*) did not significantly differ between the two feature conditions (location *M*: 0.75, *SD*: 0.18; colour *M*: 0.73, *SD*: 0.20), *t*(19) = 0.65, *p* = .524.

**Figure 2.**
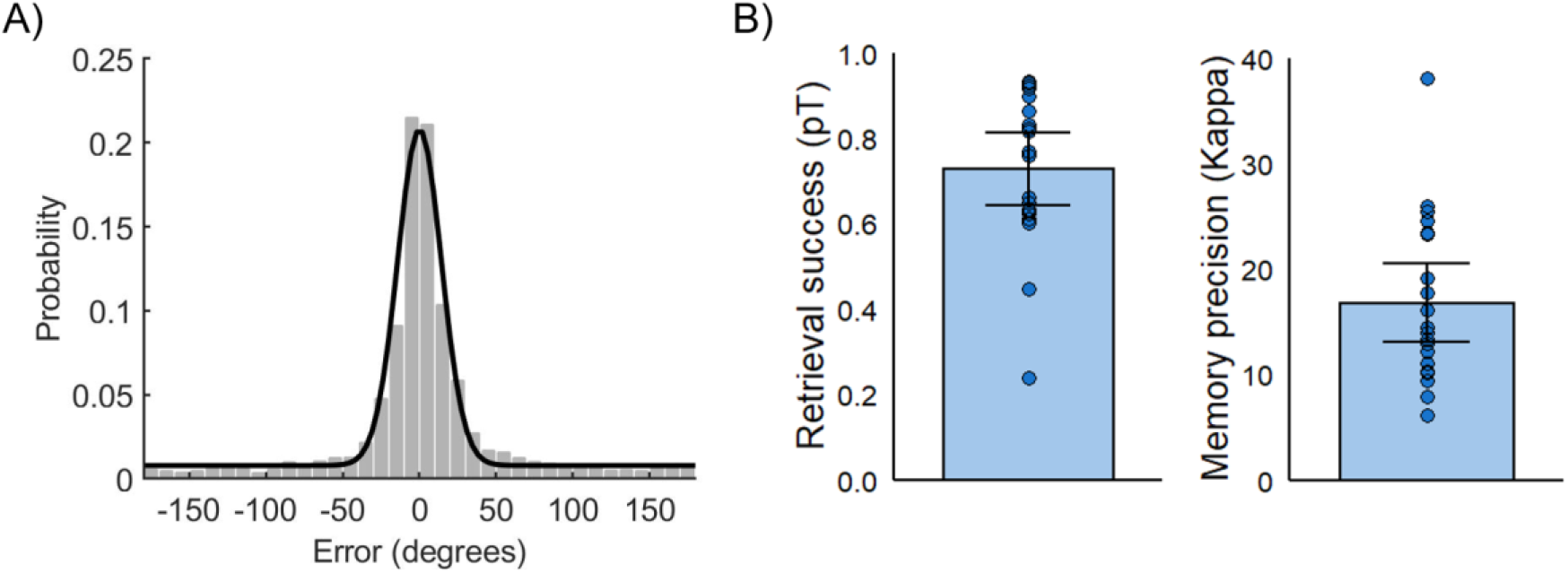
A) Distribution of retrieval errors (response – target) across all trials and participants. Black line illustrates response probabilities predicted by the two-component mixture model (von Mises + uniform distribution; model fitted to data across all participants for visualization). B) Mean model-estimated probability of successful memory retrieval (*pT*) and memory precision (*K*) across participants. Error bars display 95% confidence interval of the mean and data points individual participant parameter estimates.

Consistent with previous results (Richter et al., 2016), we also observed a moderate positive correlation between estimates of the probability of successful memory retrieval and memory precision across participants, *r*_*s*_ = .54, *p* = .014.

### fMRI results

#### Encoding activity predicting subsequent retrieval success and memory precision in a priori ROIs

Our ROI analyses focused on examining whether encoding activity in three regions typically displaying subsequent memory effects for visual information; the hippocampus, the inferior frontal gyrus and the fusiform gyrus, differentially contributes to the later success and precision of episodic memory retrieval. We first examined increases in encoding activity for trials that were subsequently successfully retrieved (absolute retrieval error ≤ 51°) in contrast to trials that were subsequently forgotten (absolute retrieval error > 51°). Within our anatomical ROIs, we observed increased encoding activity in the inferior frontal gyrus, *t*(19) = 6.75, *p* = .001, peak: −36, 27, 18, and the fusiform gyrus, *t*(19) = 8.88, *p* < .001, peak: −30, −63, −9, to predict whether object features were later successfully retrieved from memory or forgotten (see Figure 3A and 4A). In contrast, no significant subsequent retrieval success effects were detected in the hippocampus (*p*s > .151).

**Figure 3.**
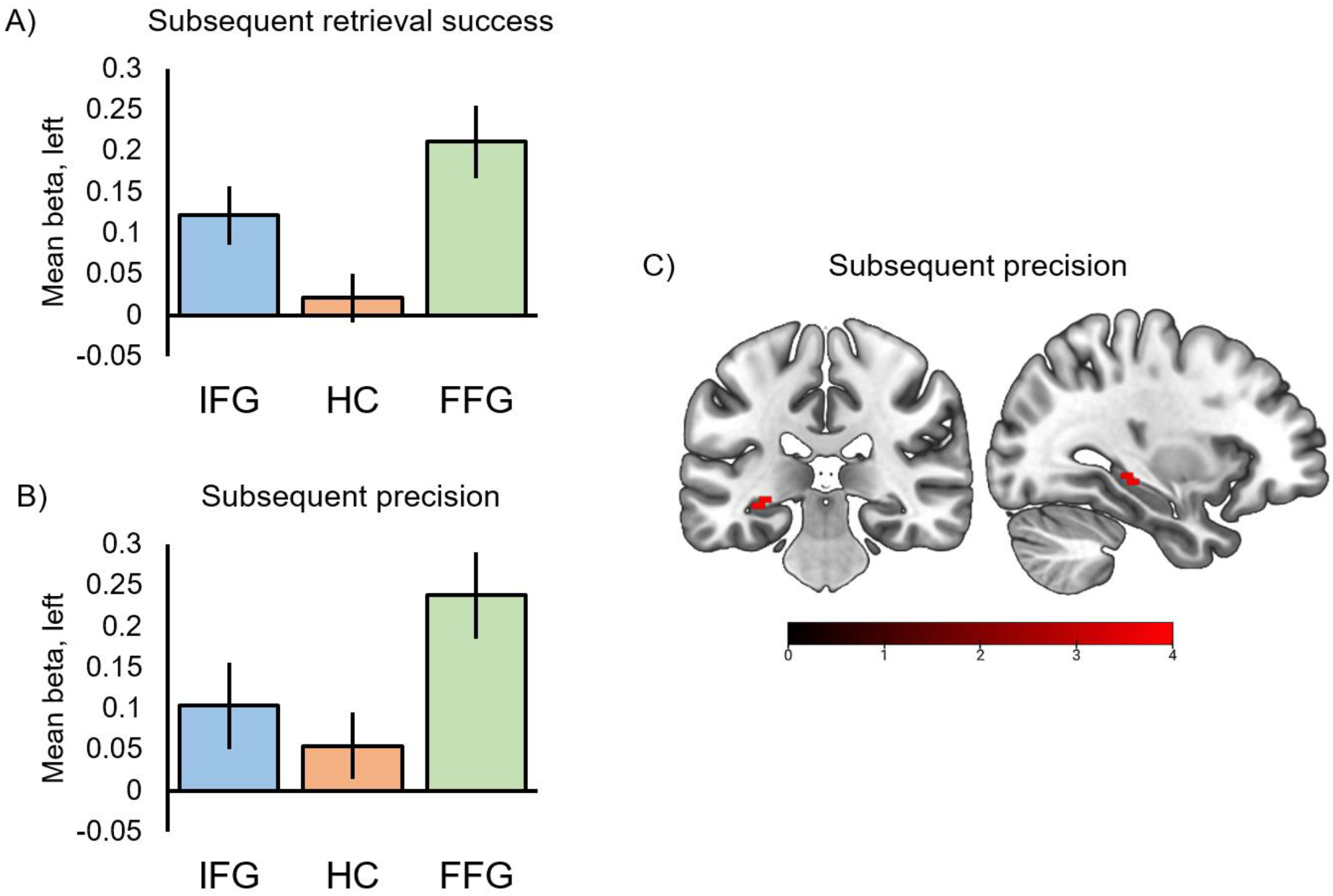
Mean parameter estimates for A) subsequent success and B) subsequent precision effects in the left inferior frontal gyrus (IFG), hippocampus (HC) and fusiform gyrus (FFG). Error bars display +/-1 SEM. C) Encoding activity correlating with the subsequent precision of memory retrieval in the hippocampal ROI (visualised at an uncorrected threshold of *p* < .001).

We next examined whether encoding activity in these regions predicted the graded precision with which object features were later successfully retrieved from memory (linear relationship between BOLD signal and precision parametric modulator). In addition to predicting which trials were successfully remembered, encoding activity in the inferior frontal gyrus, *t*(19) = 5.63, *p* = .011, peak: −57, 15, 15, and the fusiform gyrus, *t*(19) = 6.27, *p* = .001, peak: −33, −75, −18, positively correlated with the precision of later memory retrieval (see Figure 3B and 4B). Furthermore, increased encoding activity in the hippocampus, *t*(19) = 4.20, *p* = .029, peak: −33, −30, −9, was associated with greater mnemonic precision for object features (see Figure 3B and 3C). Control analyses suggested these effects to be specific to successful memory retrieval, with no significant relationships between BOLD signal and trial-by-trial variation in memory error detected across the ROIs for trials classified as forgotten (*p*s > .238).

Thus, results from the ROI analyses suggest encoding activity in the inferior frontal and fusiform cortex to support both the later success and precision of memory retrieval, while significant increases in BOLD signal in the hippocampus were observed for subsequent memory precision only. We next sought to assess whether encoding activity predicting these two aspects of later retrieval performance overlapped in any of the ROIs. Conjunction analyses indicated significant overlap between subsequent success and subsequent precision effects in both the inferior frontal *t*(19) = 4.86, *p* = .007, peak: −42, 3, 27, and the fusiform gyrus, *t*(19) = 6.12, *p* < .001, peak: −42, −57, −12, whereas no significant overlap was detected in the hippocampus (*p*s > .778). We further directly compared the mean beta-estimates for the subsequent success and subsequent precision effects in each anatomical ROI to assess evidence for disproportionate involvement in one, or the other, aspect of later retrieval performance. A within subjects ANOVA with the factors of region (HC, IFG, FFG) and memory measure (success vs. precision) did not provide evidence for a significant effect of memory measure, *F*(1, 19) = 0.05, *p* = .825, or a significant interaction between memory measure and brain region, *F*(2, 38) = 0.58, *p* = .568, however. Indeed, although no conjoint encoding activity relating to subsequent retrieval success and precision was detected in the hippocampus as described above, the mean subsequent success and subsequent precision effects did not significantly differ in the hippocampus (HC: *p* = .529), or the other regions of interest (IFG: *p* = .808, FFG: *p* = .742).

#### Encoding activity predicting subsequent retrieval success and memory precision across the whole brain

To identify any additional brain regions, beyond our *a priori* ROIs, where encoding activity predicted the later success and/or precision of memory retrieval, we further performed complementary whole brain analyses. While for subsequent memory precision, no significant clusters survived a whole brain corrected significance threshold (*p* < .050 FWE-corrected, *k* > 5; see Figure 4B for results visualized at an uncorrected threshold), activity in several regions of the dorsal and ventral visual stream, including the middle occipital gyrus, inferior parietal gyrus, fusiform gyrus, and inferior temporal gyrus, was found to predict which object features were later successfully remembered vs. forgotten (see Table 1 and Figure 4A). Whole brain conjunction analyses further indicated significant overlap between subsequent success and subsequent precision effects in the middle occipital and fusiform gyri (see Table 1 and Figure 4C).

**Table 1.** Encoding activity associated with the subsequent success and precision of memory retrieval in the whole brain analyses, p < .050 FWE, k > 5.

| **Region** | **Voxels** | **x** | **y** | **z** | **t** | **p** |
| --- | --- | --- | --- | --- | --- | --- |
| *Subsequent retrieval success* |  |  |  |  |  |  |
| L middle occipital gyrus | 355 | -36 | -87 | 12 | 10.50 | < .001 |
| L inferior parietal gyrus | 87 | -33 | -45 | 39 | 9.41 | < .001 |
| R middle occipital gyrus | 63 | 33 | -75 | 30 | 8.83 | .001 |
| R inferior temporal gyrus | 15 | 45 | -54 | -12 | 8.27 | .003 |
| R middle occipital gyrus | 21 | 42 | -81 | 9 | 7.95 | .005 |
| R fusiform gyrus | 8 | 30 | -27 | -21 | 7.33 | .016 |
| *Subsequent precision* |  |  |  |  |  |  |
| No significant clusters. |  |  |  |  |  |  |
| *Conjoint activity* |  |  |  |  |  |  |
| L middle occipital gyrus | 30 | -24 | -75 | 30 | 6.59 | .002 |
| R middle occipital gyrus | 41 | 33 | -78 | 24 | 6.39 | .004 |
| L fusiform gyrus | 30 | -42 | -57 | -12 | 6.12 | .008 |
| L middle occipital gyrus | 22 | -39 | -84 | 6 | 6.01 | .010 |

*Note.* L = left, R = right.

**Figure 4.**
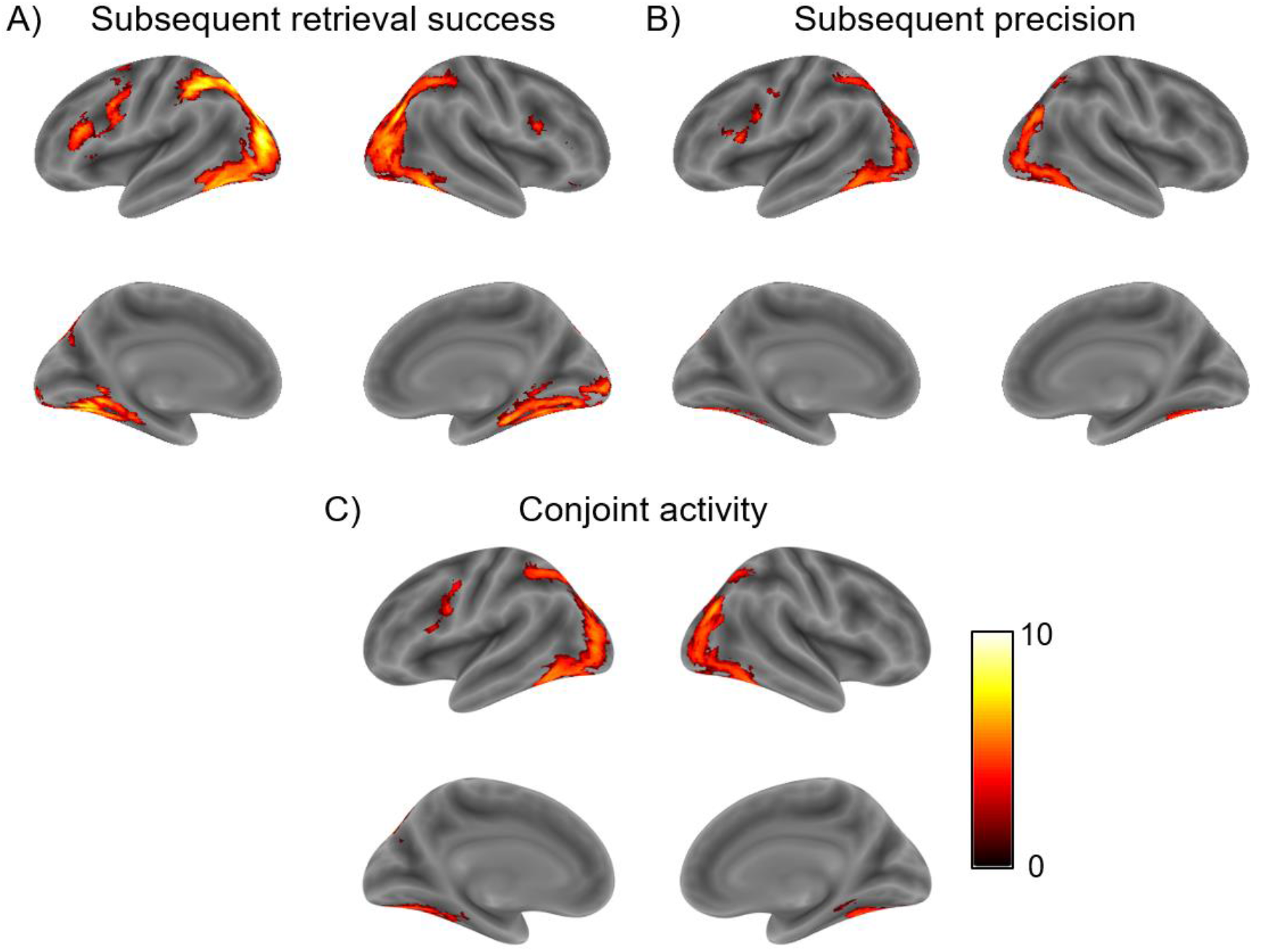
Encoding activity predicting the A) subsequent success and B) subsequent precision of memory retrieval, and C) overlap between encoding activity predicting both later success and precision of memory retrieval. Visualized at an uncorrected threshold of *p* < .001, minimum cluster of 10 voxels.

## Discussion

Amid growing interest in the neural substrates underlying the precision of episodic memory, prior studies have predominantly focused on *retrieval* processes (Cooper & Ritchey, 2019; Montchal et al., 2019; Richter et al., 2016; Stevenson et al., 2018), leaving the specific *encoding* mechanisms supporting the acquisition of high-fidelity memories largely uncharacterised. Here, we employed continuous measures of memory retrieval in combination with model-based analyses of fMRI data to segregate the encoding activity supporting the later success and precision of episodic retrieval. We observed encoding activity in overlapping cortical regions, including the inferior frontal, fusiform and middle occipital gyri, to predict both which object features were later successfully retrieved from memory versus forgotten, and the precision with which they were reconstructed. In contrast, encoding activity in the hippocampus significantly predicted the precision of later memory retrieval only. Together, these findings highlight a hippocampal-cortical basis for the formation of high-fidelity memories of perceptual information, and provide novel insight into the encoding substrates supporting the accessibility and precision of episodic memory.

The current findings demonstrating a relationship between hippocampal encoding activity and later memory precision are consistent with previous accounts emphasizing a critical role for this region in supporting detailed episodic memories (Moscovitch et al., 2016; Robin & Moscovitch, 2017). Prior neuroimaging studies have found hippocampal encoding activity to correlate with both behavioural and neural indices of later retrieval quality, including the number of event details remembered (Cooper & Ritchey, 2020; Qin et al., 2011; Staresina & Davachi, 2008), and the specificity of subsequent neural reinstatement of mnemonic content (Danker et al., 2016; Wing et al., 2015). Aligning with these accounts, we here provide evidence to demonstrate a relationship between hippocampal encoding activity and subsequent behavioural memory precision. Specifically, we observed linear increases in BOLD signal in the left hippocampus to predict the graded precision with which object features were later reconstructed from memory. Interestingly, the peak of this subsequent precision effect was located in the posterior hippocampus, consistent with emerging evidence suggesting posterior hippocampus to be particularly important for supporting the fine-grained representation of perceptual details (Brunec et al., 2018; Poppenk et al., 2013).

In contrast, we did not observe significant subsequent retrieval success effects in the hippocampus in our current paradigm, nor any significant overlap in this region between encoding activity associated with the subsequent success and subsequent precision of episodic recollection. The lack of significant retrieval success effects in the hippocampus may seem surprising given previous evidence for hippocampal encoding increases for successful versus unsuccessful encoding of associative information (Davachi, 2006; Staresina & Davachi, 2008), but aligns with the proposed involvement of the hippocampus in supporting complex, high-resolution bindings between separate event features (Yonelinas, 2013). Specifically, hippocampal involvement is postulated to increase with greater demands for representation of both high-dimensional and high-fidelity associations (Yonelinas, 2013), a proposal consistent with our current finding of greater hippocampal encoding activity with higher precision of object-feature bindings. Our findings are further in line with patient evidence demonstrating medial temporal lesions to impair both short and long term memory for high-fidelity, but not coarse-grained, bindings (Koen et al., 2017; Nilakantan et al., 2018), and suggest a potential role for a deficient hippocampal encoding mechanism in such impairments. However, it is important to note that despite a lack of significant subsequent retrieval success effects in the hippocampus, we did not observe evidence for significantly greater involvement of hippocampal encoding activity in subsequent memory precision when compared to subsequent retrieval success either.

Interestingly, a prior study employing a similar paradigm to the one used here found hippocampal *retrieval* activity to be associated with the success, but not precision, of episodic memory retrieval (Richter et al., 2016). This apparent difference between encoding and retrieval effects in the hippocampus may reflect differential involvement of hippocampal *pattern separation* and *pattern completion* during encoding and retrieval, respectively. Specifically, at encoding, hippocampal pattern separation is thought to facilitate the formation of distinct memory traces (Norman & O’Reilly, 2003; Yassa & Stark, 2011), thus posing a putative prerequisite for later high-fidelity memory retrieval (Moscovitch et al., 2016; Xie et al., 2020). In contrast, at retrieval, hippocampal *pattern completion* enables access to stored memory representations when presented with a noisy or partial cue (Norman, 2010; Norman & O’Reilly, 2003), while retrieval precision may be more reliant on cortical regions including the lateral parietal cortex (Richter et al., 2016; Ritchey & Cooper, 2020). Pattern separation and pattern completion are thought to be supported by different subfields of the hippocampus (Hunsaker & Kesner, 2013). An interesting question for future studies to explore is whether different hippocampal subfields also support the precision and accessibility of episodic memory during encoding and retrieval, respectively.

Beyond the hippocampus, we observed activity in overlapping cortical regions, including the inferior frontal, fusiform, and middle occipital gyrus, to predict both the later success and precision of episodic memory retrieval. Our finding of left inferior frontal involvement in subsequent retrieval success and precision is consistent with previous evidence implicating this region in cognitive control of memory encoding, supporting successful memory formation across a range of encoding tasks and mnemonic content (Blumenfeld et al., 2011; Blumenfeld & Ranganath, 2006; Murray & Ranganath, 2007; Park & Rugg, 2008, 2011). Specifically, ventrolateral regions of the prefrontal cortex have been proposed to support the attentional selection and elaboration of goal-relevant information, leading to formation of strong and distinctive memory traces for specific item features (Blumenfeld et al., 2014; Blumenfeld & Ranganath, 2007; Simons & Spiers, 2003). Such selective encoding processes supported by this region may act to enhance the representation of goal-relevant features in posterior perceptual regions (Chun & Turk-Browne, 2007; Gilbert & Li, 2013; Sprague et al., 2015; Xue et al., 2013), and/or modulate hippocampal encoding more directly (Aly & Turk-Browne, 2017; Carr et al., 2013), aiding the formation of durable and precise memory representations.

The current results further emphasize the role of perceptual regions in supporting the formation of accessible and precise memory traces. Specifically, we observed encoding activity in the fusiform gyrus, a region typically associated with object perception and memory (Bar et al., 2001; Haxby et al., 2001; Vaidya et al., 2002), to predict both the later success and precision of object feature retrieval. This finding is consistent with previous studies that have observed memory-related activity increases in the fusiform gyrus during episodic encoding (reviewed in Kim, 2011; Spaniol et al., 2009), potentially playing an important role in formation of detailed object representations (Garoff et al., 2005; Kensinger et al., 2007), and with evidence suggesting representational specificity in the occipitotemporal cortex during encoding to predict subsequent memory performance (Gordon et al., 2014; Ward et al., 2013; Xue et al., 2010). Beyond our regions of interest, we further observed that encoding activity in a wider network of ventral and dorsal visual regions predicted the subsequent success of memory retrieval. Of these regions, conjoint subsequent retrieval success and precision effects were observed in the middle occipital gyrus. The involvement of a broader set of ventral and dorsal visual regions, implicated in processing of “what” and “where” visual information, respectively (Haxby et al., 2001; Mishkin et al., 1983), may reflect the demands of the current task for processing both item and spatial stimulus features, such as object colour and location, respectively. This interpretation is consistent with the idea that activity in content-specific perceptual regions supports successful memory formation for distinct event attributes (Cooper & Ritchey, 2020; Gottlieb et al., 2010). At the behavioural level, we observed the probability and precision of successful memory retrieval to correlate moderately across participants. Although both behavioural (Harlow & Donaldson, 2013; Harlow & Yonelinas, 2016, but see Schurgin et al., 2020) and neural (Richter et al., 2016) evidence suggests partial independence, these two aspects of episodic retrieval are also typically characterised by a degree of shared variance (Richter et al., 2016). Our current findings highlight common cortical encoding substrates as one possible factor contributing to this shared variance. Notably, the representation of stimulus-specific information in posterior perceptual regions, and strategic encoding operations mediated by the lateral prefrontal cortex, may be important for the formation of durable memory traces that are readily accessible from memory and can be reconstructed with a high degree of precision. The computational properties of the hippocampus, on the other hand, may render this structure important for the acquisition of differentiated, high-fidelity, mnemonic representations (LaRocque et al., 2013; Norman & O’Reilly, 2003; Wing et al., 2020).

In summary, the current study aimed to elucidate the encoding mechanisms supporting the formation of accessible and precise memory traces. We observed encoding activity in prefrontal and posterior perceptual regions to support both the later success and precision of episodic memory retrieval. In contrast, activity in the hippocampus was found to significantly predict later memory precision only. These results highlight the importance of hippocampal-cortical encoding mechanisms for the formation of precise episodic memories, and have important implications for understanding memory deficits in distinct populations.

## Acknowledgements

This study was funded by BBSRC grant BB/L02263X/1 and James S. McDonnell Foundation Scholar Award #220020333, and was carried out within the University of Cambridge Behavioural and Clinical Neuroscience Institute, funded by a joint award from the Medical Research Council and the Wellcome Trust. We are grateful to Paul Bays for valuable advice and to the staff of the MRC Cognition and Brain Sciences Unit MRI facility for scanning assistance.

## Notes

### Competing Interest Statement

The authors have declared no competing interest.

